# Anticipatory distractor suppression elicited by statistical regularities in visual search

**DOI:** 10.1101/599134

**Authors:** Benchi Wang, Joram van Driel, Eduard Ort, Jan Theeuwes

**Affiliations:** Department of Experimental and Applied Psychology, Vrije Universiteit Amsterdam, Amsterdam, the Netherlands

**Keywords:** Attentional capture, statistical learning, suppression, alpha enhancement

## Abstract

Salient yet irrelevant objects often capture our attention and interfere with our daily tasks. Distraction by salient objects can be reduced by suppressing the location where they are likely to appear. The question we addressed here was whether suppression of frequent distractors is already implemented beforehand, in anticipation of the stimulus. Using electroencephalography (EEG), we recorded cortical activity of human participants searching for a target while ignoring a salient distractor. The distractor was presented more often at one location than at any other location. We found reduced capture for distractors at frequent locations, indicating that participants learned to avoid distraction. Critically, we found evidence for *proactive suppression* as already *prior to display onset*, there was enhanced power in parieto-occipital alpha oscillations contralateral to the frequent distractor location – a signal known to occur in anticipation of irrelevant information. Locked to display onset, event-related potentials analysis showed a distractor-suppression-related P_D_ component for this location. Importantly, this P_D_ was found regardless of whether distracting information was presented at the frequent location. In addition, there was an early P_D_ component representing an early attentional index of the frequent distractor location. Our results are show anticipatory (proactive) suppression of frequent distractor locations in visual search already starting prior to display onset.

## Introduction

In everyday life, we try to attend to events that are relevant to us and ignore information that could distract us. For example, when driving along a busy road, we devote attention to relevant objects such road signs and other cars, while ignoring salient irrelevant objects such as billboards. The extent to which we are able to avoid distraction from salient events has been a central question for many years.

Traditionally, attentional selection is assumed to be the result of the interaction between the goals of the observer (current selection goals) and the physical properties of the visual environment (salience of the objects; Egeth & Yantis, 1997; Itti & Koch, 2001; Theeuwes, 2010). Recently, it is pointed out that in many cases selection is neither the result of goals of the observer nor the result of stimulus-driven (i.e., bottom-up saliency) factors (Awh, Belopolsky, & Theeuwes, 2012; Failing & Theeuwes, 2017; Theeuwes, 2018). A third category labelled as “selection history” referred to those instances in which selection is the result of previous attentional deployments (Theeuwes, 2019). These previous deployments can result in lingering and enduring selection biases that are unrelated to the current goals nor related to the stimulus-driven saliency of objects.

A well-known example of how selection history affects attentional selection is research known as “contextual cueing” demonstrating that search for a target is facilitated when it appears in a visual lay-out that was previously searched relative to visual lay-outs that were never seen before (Chun & Jiang, 1998, 1999; Jiang & Chun, 2001). These studies have highlighted that people pick up on the statistical regularities present in the display, such that they selectively attend those locations that are relevant for the task. Furthermore, recently, it was demonstrated that people cannot only activate locations that are relevant but also selectively inhibit salient distractors that may distract them (Failing, Wang, & Theeuwes, 2019; Wang & Theeuwes, 2018a, 2018b, 2018c). In these studies, Wang and Theeuwes used the well-established additional singleton task (Theeuwes, 1992) in which observers search for a unique shape singleton while a unique and salient color singleton is simultaneously present. Because observers cannot search for one specific target feature, large capture effects are expected (see Wang & Theeuwes, 2018c). In this paradigm we manipulated the distributional properties of the location of salient distractor. It was shown that when a salient distractor appeared more often in one location than in all other locations, its distracting effect (the extent to which it captured attention) was attenuated. Crucially, there was a spatial gradient of suppression around this location, suggesting that this frequent distractor location competed less for attention than all other locations. They also showed that when the target happened to be presented at that location, its selection was hampered (see Ferrante et al., 2018 for similar results).

The finding that we were able to inhibit frequent distractor locations that were likely to contain salient distractor has important implications for theories of visual selection and other related fields. Event-related potential (ERP) studies have shown that, ignoring of a salient distractor is typically accompanied by a distractor positivity (P_D_) component in the ERP waveform (Burra & Kerzel, 2014; Feldmann-Wüstefeld, Uengoer, & Schubö, 2015; Gaspar & McDonald, 2014; Sawaki & Luck, 2010), indicating spatial suppression of its location during search (Hickey, Di Lollo, & McDonald, 2008). However, it remains unclear whether such suppression of frequent distractor locations is already implemented beforehand, in anticipation of the stimulus.

In the current study, using electroencephalography (EEG), we recorded cortical activity of human participants who were searching for a target, while ignoring a salient distractor. The salient distractor was presented more often in one location (high-probability location) than in all other locations (low-probability location). Consistent with a suppression account and our previous behavioral studies (Wang & Theeuwes, 2018a, 2018b, 2018c), we expected to observe the classic distractor-related P_D_ component in the ERP signal when a distractor was presented at the high-probability location. However, if spatial suppression to this location is applied well before display onset, we also expected to observe increased power in parieto-occipital alpha oscillations contralateral to the location that was more likely to contain the distractor relative to the ipsilateral location (Worden, Foxe, Wang, & Simpson, 2000; see Klimesch, 1999 for a review). We also conducted a baseline experiment in which all locations were equally likely to contain a distractor (i.e., without any statistical regularities), and here we expected to see no EEG components related to spatial suppression.

## Method

The study was approved by the Ethical Review Committee of the Faculty of Behavioral and Movement Sciences of the Vrije Universiteit Amsterdam.

### Participants

Separate groups with each including 18 participants (13 women in Experiment 1 with a mean age of 20.9 years; 17 women in Experiment 2 with a mean age of 19.7 years) took part in the study. All participants provided written informed consent prior to the experiment and were reimbursed for their participation either with money or course credit. They reported normal color vision and normal or corrected-to-normal visual acuity.

### Apparatus and stimuli

Stimulus presentation and response registration were controlled by custom scripts written in Python run on an HP Compaq 8000 Elite computer with a 22-in. liquid crystal display (LCD) color monitor (Samsung Syncmaster 2233, 1680 × 1050 pixels at 120 Hz). Participants were seated in a sound-attenuated and dimly lit laboratory, at a viewing distance to the monitor of 75 cm. They were instructed to sit as relaxed as possible to minimize muscle activity and other electrophysiological noise to appear in the Electroencephalogram (EEG) signal.

The primary experimental display contained a visual search array consisting of eight discrete stimuli with different shapes (one circle vs. seven unfilled diamonds, or vice versa), with each shape contained a vertical or horizontal grey line (0.3° × 1.5°) inside. These stimuli were presented on an imaginary circle with a radius of 4°, centered at the fixation (a white cross measuring 1° × 1°), against a black background (6 cd/m^2^). The circle’s radius was 1°, the unfilled diamond was subtended 2° × 2°, and each shape had a red or green outline (the color was chosen from CIE Lab color space, centered at L = 70, a = 5, b = 0, with a radius of 60, with an almost equal luminance range: 36-44 cd/m^2^).

### Procedure and design

A fixation cross appeared and remained visible throughout the trial. After 1350 ms the search array was presented for 2450 ms or until the participant responded. Participants had to search for a circle (target) among seven diamonds (distractors), or *vice versa*, and indicate whether the line segment inside the target was vertical or horizontal, by pressing the *up* or *left* key on the keyboard as fast as possible. Because the target singleton could either be a diamond between circles or a circle between diamonds, observers most likely adopted a “singleton detection mode” which makes it impossible to selectively attend one specific feature (see Bacon & Egeth, 1994). The inter-trial interval (ITI) was between 0 and 350 ms at random.

### Experiment 1: with high-probability location

The target was presented in each trial, and it was equally likely to be a circle or a diamond. A uniquely colored distractor singleton appeared in 66% of the trials, with the same shape as other distractors but a different color (red or green with an equal probability). All conditions were randomized within blocks. Those display-elements were presented at eight locations from the imaginary ring with a 4° radius. The target and distractor singleton were only presented along the vertical or horizontal axis. The remaining four locations served as neutral display-elements. For each participant, one of the horizontal locations contained a distractor singleton in 55% of trials (high-probability location); for other three locations (on the cardinal axis) the probability of containing a distractor singleton was 15% each (low-probability location). The high-probability location was counterbalanced across participants. In the distractor singleton present condition, the target appeared equally often at each possible location, except for the high-probability location^1^. In the no-distractor condition, the target appeared equally often at each possible location.

Participants completed 40 practice trials and 13 blocks, with each containing 120 trials. Warning messages were presented if they did not respond, or pressed the wrong key. Depending on the singleton distractors location and target location, in total five conditions were analyzed: 1) the distractor singleton was presented at the high-probability location while the target was presented vertically; 2) the target was presented at high-probability location while the distractor singleton was either absent or presented vertically; 3) the distractor singleton was either absent or presented vertically, while the target was presented vertically; 4) the distractor singleton was presented along the horizontal axis at the lateral low-probability location while the target was presented vertically; 5) the distractor singleton was either absent or presented vertically, while the target was presented along the horizontal axis at the lateral low-probability location.

### Experiment 2: without high-probability location

Experiment 2 served as a baseline experiment to compare it with the results acquired in Experiment 1. In Experiment 2, we used the same design as in Experiment 1 except that here, all four potential target and singleton distractor locations were equiprobable. Participants completed 40 practice trials and 9 blocks each containing 120 trials. As there was no high-probability location, only three conditions were investigated: 1) the distractor singleton was presented laterally while the target was presented vertically; 2) the target was presented laterally when the distractor singleton was absent or presented vertically; 3) the distractor singleton was either absent or presented vertically, while the target was also presented vertically. Note that, in both experiments, both target and distractor singleton were salient objects.

### Data recording and preprocessing

Electroencephalographic (EEG) data were collected using a BioSemi ActiveTwo system (www.biosemi.com) from 64 electrodes, placed according to the extended 10–20 system and digitized at 512 Hz. Additionally, to record eye movements, the vertical electro-oculogram (VEOG) was recorded from electrodes located 2 cm above and below the right eye, and the horizontal electro-oculogram (HEOG) was recorded from electrodes 1 cm lateral to the external canthi.

EEG data were re-referenced to the mean of left and right earlobes and high-pass filtered using a cut-off of 1.5 Hz (for independent component analysis [ICA] only) and 0.1 Hz (for final analyses). High-pass filtering was done with default settings of the pop_basicfilter function of ERPLAB (Lopez-Calderon & Luck, 2014), using a Butterworth IIR filter. Continuous EEG was first epoched from –3000 to 3500 ms relative to search display onset (to avoid edge artifacts resulting from wavelet convolution; see below). Trial rejection and ICA were done on smaller epochs (–1500 to 1500 ms) to avoid overlapping time points across epochs. To improve ICA, epochs were baseline-normalized using the whole epoch as baseline (Groppe, Makeig, & Kutas, 2009).

Malfunctioning electrodes were visually detected and temporarily removed from the data. An adapted version of an automatic trial-rejection procedure as implemented in the Fieldtrip toolbox (Oostenveld, Fries, Maris, & Schoffelen, 2011) was used to detect Electromygram (EMG) artifacts. Specifically, we used a 110 –140 Hz bandpass filter to capture muscle activity, and allowed for variable z-score cutoffs per subject based on the within-subject variance of z-scores. After trial rejection, ICA as implemented in the EEGLAB toolbox (Delorme & Makeig, 2004) was performed on the clean electrodes only. Together with the VEOG and HEOG signals, we visually inspected and removed ICA components that captured eye-blinks, eye-movement, or other artifacts that were clearly not brain-driven. Afterwards, we interpolated the malfunctioning electrodes identified earlier using spherical spline interpolation as implemented in EEGLAB’s eeg_interp.m function. Trials with an incorrect (or no) response were also excluded from further analyses. All steps together left 87.4% (± 6.0% across subjects) trials in Experiment 1, and 73.5% (± 3.4% across subjects) trails in Experiment 2.

## Time-frequency analysis

### Time-frequency decomposition: Laplacian

Before time-frequency decomposition, we estimated the surface Laplacian of the EEG data (Perrin, Pernier, Bertrand, & Echallier, 1989), which is equivalent to the current source density approach. This method acts as a spatial high-pass filter by accentuating local effects while filtering out distant effects due to volume conduction; thus, it sharpens the EEG topography (Cohen, 2014; Kayser & Tenke, 2015). For estimating the surface Laplacian, we used a 10th-order Legendre polynomial, and *λ* was set at 10^−5^.

### Morlet wavelet convolution

The epoched EEG time series were decomposed into their time-frequency representations using custom-written MATLAB scripts (github.com/joramvd/tfdecomp). The time series were convolved with a set of Morlet wavelets with frequencies ranging from 1 to 40 Hz in 25 logarithmically spaced steps. The complex wavelets were created by multiplying sine waves (*e*^*i*2π*ft*^ where *i* is the complex operator, *f* is frequency, and *t* is time) with a Gaussian (*e*^*−t*^2*/* 2*s*^2^, where *s* is the width of the Gaussian). The width of the Gaussian was set as *s* = *δ/*2π*f*, where *δ* represents the number of cycles of each wavelet, logarithmically spaced between 3 and 12 to have a good trade-off between temporal and frequency precision. Convolution was applied in the frequency domain: The Fast Fourier Transform was applied to both the EEG data and the Morlet wavelets, and these were multiplied in the frequency domain, after which the inverse Fast Fourier Transform was applied. The frequency-specific power at each time point was defined as the squared magnitude of the complex signal resulting from the convolution (real(Z_t_)^2^ + imag(Z_t_)^2^). Raw power was transformed into one common scale across frequencies using a within-subject permutation testing procedure (see below).

### Frequency band and time window of interest

Based on previous studies, we selected PO3/4, PO7/8, and O1/2 for the analysis (van Driel, Gunseli, Meeter, & Olivers, 2017; Vries, van Driel, & Olivers, 2017). We were specifically interested in the lower (7.5–10 Hz) and upper (10–12.5 Hz) alpha-frequency ranges (see Klimesch, 1996, 1999, 2012 for reviews), in order to track how attention is guided by implicit regularities before the search display onset, in a time window comprising the inter-trial interval (–1500 to 0 ms). Moreover, because participants could only anticipate the high-probability distractor location, and not the target location nor the actual distractor location, we combined all five conditions together in both experiments for the pre-stimulus alpha-band analysis.

### Z-transformation of lateralized power

Raw power can only take positive values and has a 1/f power scaling, which requires a transformation such as dB conversion or percent signal change (Cohen, 2014). Such relative metrics, in turn, require a baseline time range, for which typically a pre-stimulus interval is chosen. However, for this analysis we were specifically interested in a between-group comparison of condition-average lateralized (contra-versus ipsilateral to anticipated distractor location) alpha power during this pre-stimulus interval. We thus Z-transformed time-frequency power through a within-subject statistical procedure. For each contra-and ipsilateral channel-pair (thus excluding midline electrodes), and each time-frequency point, we first computed the trial-average difference-scores of contra-minus ipsilateral power. Next, at each of 1000 permutations, we swapped for a random subset of trials the order of this comparison (i.e. ipsi-minus contralateral), and computed the trial-average across all trials. This gave a distribution of “lateralized” power values under the null hypothesis of no difference between contra-and ipsilateral channels. The observed lateralized power values were then statistically transformed with respect to the mean and standard deviation of this distribution, by computing their Z-values. This procedure yields a metric that shows interpretable dynamics over time as well as frequency. Since there was no anticipated distractor location in the baseline experiment (Exp. 2), we labeled one lateral side as “contralateral” and the other side as “ipsilateral” consistently for all participants and accordingly computed the same Z-transformation for time-frequency power as was done for the real testing (Exp. 1) with predictable lateralized distractors.

### Statistics

At the group level, time-frequency maps of power of the selected electrode groups were first statistically tested against baseline within each experimental group, using a permutation-based approach with cluster-size correction. However, with permutation testing, the more points are added to the analysis, the less sensitive it becomes. If it is known in advance that a specific frequency band is likely to exhibit the effect of interest, then one should incorporate this prior knowledge and limit the statistical test to this certain band (Maris & Oostenveld, 2007; Oostenveld et al., 2011). Our a priori hypothesis pertained specifically to pre-onset alpha band enhancement; thus, in addition to a group-specific descriptive test over all time-frequency points, we further narrowed our hypothesis-test of a group difference, to the alpha-band only.

We ran two types of cluster-corrected permutation tests: 1) within each group, we used one-sample t-tests across time and frequency; at each test, the sign of the power values was shuffled across subjects in 2000 permutations to yield a null-distribution against which observed clusters of summed t-values were thresholded using *p* < 0.05; 2) between groups, we used independent-samples t-tests, comparing the two groups across time and frequency, and on alpha-band average power over time; here, we shuffled participants across experiment in 2000 permutations for the null distribution.

### Event-related potential (ERP) analysis

ERP analysis was done on a pre-specified time window from –100 ms to 900 ms relative to display onset. To obtain ERPs, single-trial EEG data were first baseline-corrected using a pre-stimulus baseline period from –100 to 0 ms, after which data were trial-averaged.

### Experiment 1: with high-probability location

For each participant, mean contralateral and ipsilateral activity over conditions relative to the high-probability location was calculated for electrodes PO7/8. To determine the time window used for further statistical analyses of different ERP components, the lateralized ERP (contralateral minus ipsilateral to the high-probability location) was calculated, showing lateralized activity due to the high-probability distractor location; and the time window was chosen also based on previous studies (Eimer, 1996; Feldmann-Wüstefeld & Vogel, 2018; Hickey et al., 2008; Hickey, McDonald, & Theeuwes, 2006). Thus, P1 epoch was determined as ± 20 ms from the most positive peak between 50 ms and 150 ms. The P_D_ epoch was determined as ± 20 ms from the most positive peak between 150 ms and 250 ms. When the target was presented at the lateral low-probability location, the target N2pc epoch was determined as ± 20 ms from the most *positive* peak between 150 ms and 250 ms (note that the positive peak was used here, because the lateralized ERP was defined with respect to the high-probability distractor location). When the target was presented at the high-probability location, the target-N2pc epoch was determined as the most *negative* peak between 250 ms and 350 ms, because we assumed a delayed target selection due to the suppression on the high-probability location. The mean amplitude for these time windows was calculated separately for laterality (*contra* vs. *ipsi*), each condition, and each participant.

### Experiment 2: without high-probability location

For each participant, mean contralateral and ipsilateral activity over conditions relative to the target or distractor singleton was calculated for electrodes PO7/8. To determine the time window used for further statistical analyses of different ERP components, both the lateralized ERP (contralateral minus ipsilateral) for trials with lateral target and for trials with lateral distractor singleton were collapsed, showing lateralized activity due to targets or distractor singletons. The target-N2pc epoch was determined as ± 15 ms from the most *negative* peak between 200 ms and 300 ms in the lateral-targets waveform. The mean amplitude for these time windows was calculated separately for laterality (*contra* vs. *ipsi*), each condition, and each participant.

## Results

### Behavior

Consistent with Wang and Theeuwes (2018a, 2018b), distractors presented at the high-probability location interfered less (828 ms) than distractors at the low-probability location (865 ms), *t*(17) = 3.42, *p* = .003. Compared to the no-distractor condition (777 ms), there were significant attentional capture effects for a distractor present at the high-probability location, *t*(17) = 4.26, *p* = .001, and the low-probability location, *t*(17) = 8.61, *p* < .001. Also, consistent with earlier studies (Wang & Theeuwes, 2018a, 2018b, 2018c), target selection was less efficient when the target appeared at the high-probability distractor location (808 ms) relative to the low-probability distractor location (767 ms), *t*(17) = 3.73, *p* = .002. To show that these effects were due to the predictability of the high-probability location, we also ran a baseline experiment (Experiment 2) in which all locations were equally likely to contain a salient distractor singleton. Consistent with earlier studies (Theeuwes, 1991, 1992), this showed a classic attentional capture effect, with slower RTs when the distractor singleton was present (850 ms) compared to when it was absent (773 ms), *t*(17) = 10.59, *p* < .001. See *Appendix* for more details.

### Pre-stimulus: Local alpha power

As the measure of anticipation we chose lateralized alpha power (following (Klimesch, 1996). We analyzed the oscillation in the alpha band (7.5 – 12.5 Hz) for electrodes PO3/4, PO7/8, and O1/2, (see Methods for oscillation band and electrode selection) and found a lateralized effect consisting of a stronger enhancement in power for contra-versus ipsilateral relative to the high-probability location in Experiment 1 (see Figure 1A). Importantly, there was no lateralized enhancement in Experiment 2 (see Figure 1B). A permutation test across time and frequency comparing Experiments 1 and 2 confirmed this finding, showing that the significant difference (*p* < .05 [uncorrected]) between groups was mainly seen in the alpha band (see Figure 1C upper panels for an illustration). Importantly, a cluster-based permutation test on the alpha-band comparing Experiments 1 and 2 revealed a reliable difference between the two groups in the -1220 and -660 ms interval prior to display onset (*p* < .05; see Figure 1C lower left panel for an illustration). Also, even without using the cluster-based permutation test, when comparing the average increased alpha power between the two groups, there was a reliable difference before display onset, *t* = 2.37, *p* = .03. We interpret these results as reflecting anticipatory suppression of the high-probability location.

**Figure 1.**
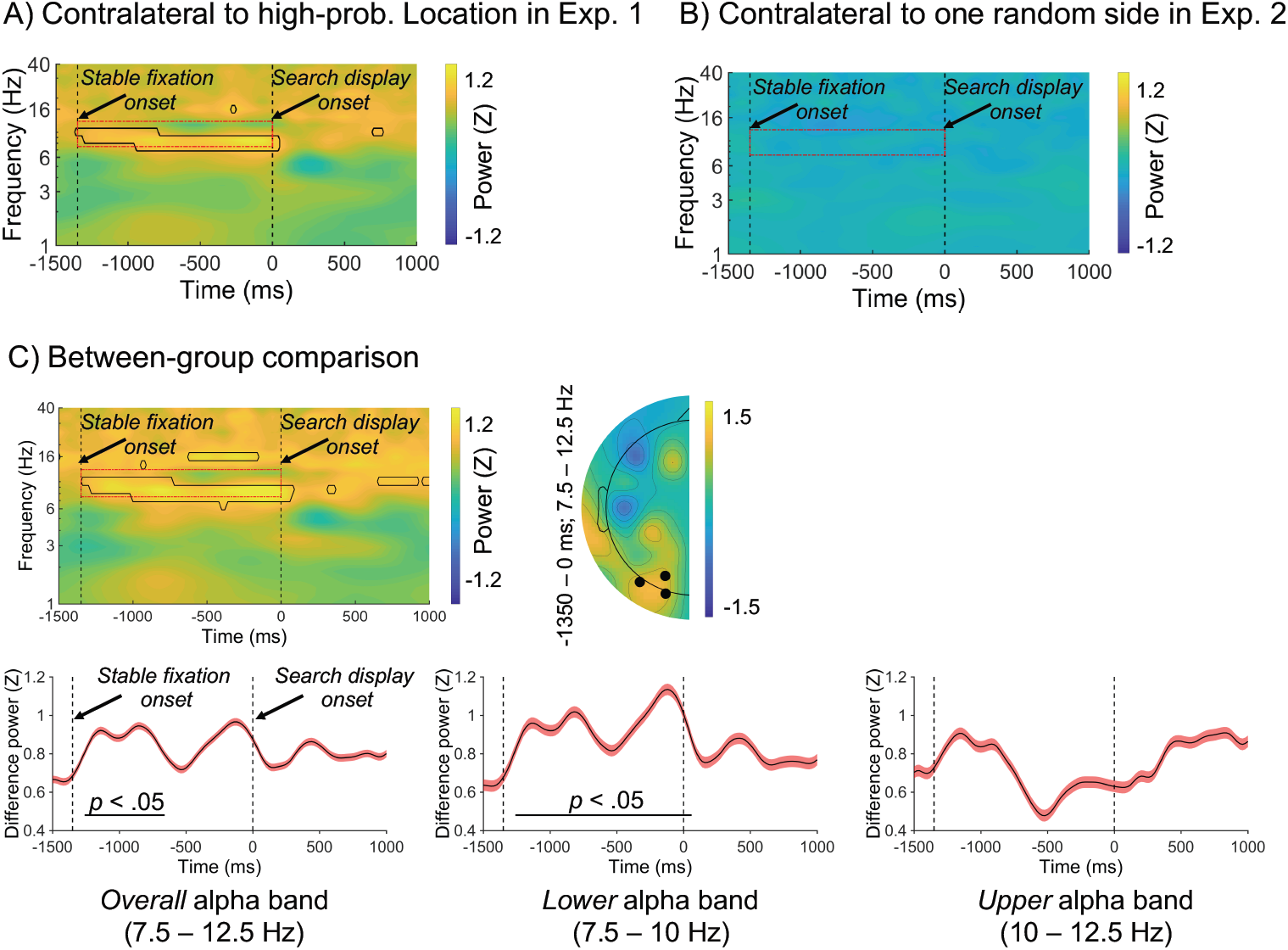
Lateralized posterior alpha power in Experiment 1 with high-probability location. (A), lateralized posterior alpha power in Experiment 2 as a baseline (B), and their comparison results (C). Sub-figures A and B show time-frequency plots of contralateral minus ipsilateral power at the average of O1/2, PO3/4, and PO7/8 in different experiments (note: red dashed outline indicates the alpha band oscillation between stable fixation and search display onset, and black solid outline indicates [uncorrected] significant power difference between contralateral and ipsilateral in grand average at *p* < .05). In sub-figure-C, upper left panel shows time-frequency plots of Experiment 1 minus Experiment 2 (baseline) power at the average of O1/2, PO3/4, and PO7/8; upper right panel shows half topographies, grand average scalp distributions of alpha power at ipsilateral electrodes subtracted from those at contralateral electrodes, collapsed across hemispheres, and power is averaged over the alpha band oscillation between stable fixation and search display onset; lower panels present time series of contralateral minus ipsilateral alpha power difference between Experiments 1 and 2 for overall alpha band, lower alpha band, and upper alpha band (note: black horizontal line represents time points with a significant difference between experiments after cluster-size thresholding at *p* < .05). Red shadow area means ± SEM of difference values.

Previous research has shown that lower alpha band (7.5-10 Hz) is more related to short-lasting expectation and attentional processing, while the upper alpha band (10-12.5 Hz) is more related to semantic memory processing (see Klimesch, 1996, 1999, 2012 for reviews). Given these findings, we divided the alpha band in two to explore whether the anticipatory suppression effect related to the high-probability location would indeed only occur in the lower alpha band. For the lower alpha band, a cluster-based permutation test between experiments showed that the power enhancement was significant from -1260 to 60 ms relative to display onset (*p* < .05, see Figure 1C middle panel). No effects were observed for the upper alpha band (see Figure 1C right panel).

The current results suggest that observers start suppressing the high-probability location about ∼1200 ms before display onset. Moreover, this pre-stimulus suppression (i.e., implicit expectation that a distractor would appear on the high-probability location) was mainly reflected in the lower alpha band, which is consistent with previous studies by Klimesch (see Klimesch, 2012 for a review).

### Post-stimulus: Distractor Positivity (P_D_)

To provide converging evidence that the high-probability location is suppressed, we first examined whether a P_D_ was elicited when the distractor singleton was presented at the high-versus the low-probability location, following earlier work linking the P_D_ component to distractor suppression (Hickey et al., 2008). Consistent with the behavioral data showing reduced capture, we observed a P_D_ when the distractor was presented in the high probability location, and not when it was presented in the low-probability condition. A two-way ANOVA with within-subjects factors *laterality* (contra-vs. ipsilateral to the distractor singleton) and *distractor location* (high-vs. low-probability) was conducted on the average ERP amplitude within the P_D_ time window (186 ms – 226 ms; see Methods section for choice of time window). The results showed a main effect for laterality, *F*(1,17) = 9.51, *p* = .007, *η* _*p*_ ^2^ = .36, but not for distractor location, *F*(1,17) = 2.92, *p* = .105, *η* _*p*_ ^2^ = .15. However, importantly, the interaction was reliable, *F*(1,17) = 4.98, *p* = .039, *η* _*p*_ ^2^ = .23. Planned comparisons tracing the source of the interaction indeed showed a reliable P_D_ only for high-probability locations, *t*(17) = 3.15, *p* = .006 (see Figure 2 upper middle and lower left panels)^2^.

**Figure 2.**
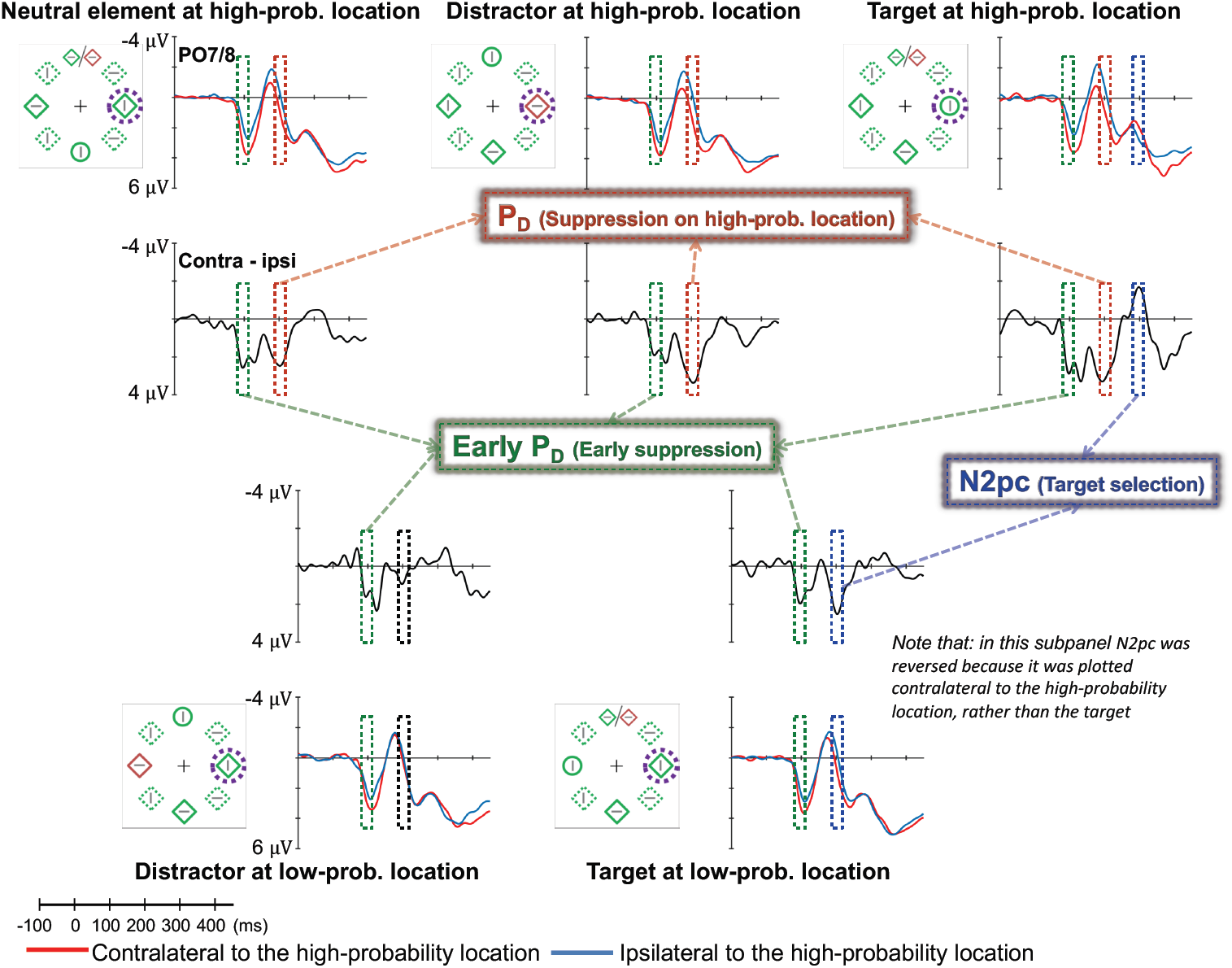
Examples of the search displays in various conditions in Experiment 1. The dashed circles indicate the high-probability location. The display-elements with solid outline indicate possible target and distractor singleton locations; the display-elements with dashed outline indicate locations that never contained a target or distractor singleton. Each sub-panel shows, per condition, a grand average ERP recorded at PO7/PO8 and the lateralized difference waves (contra-minus ipsilateral). Green, red, and blue dashed squares indicate the time window that was used to calculate mean amplitude of P1, P_D_, and N2pc, respectively. *Note that, in the lower right panel, the ERP waves were plotted contralateral to the high-probability location, but the target was presented at the low-probability location. Thus, the N2pc was reversed here, suggesting a selection towards the target.* All waveforms were low-pass filtered at 30 Hz (half power cutoff, 24 dB) for visualization.

One possible reason for not observing a P_D_ for distractors presented at the low-probability location, might be that there are opposing lateralized suppression effects working at the same time (i.e., the spatial suppression of the high-probability location and the feature suppression of the distractor singleton at the opposing low probability location). If the P_D_ reflecting the suppression on high-probability location is larger than the P_D_ reflecting the suppression on the distractor at the low probability location than it is possible that this larger P_D_ overshadows that the smaller distractor generated P_D._ Note however, it is also entirely possible that the distractor at the low probability location does not generate a P_D_ at all, as the distractor at this location generate a strong capture effect.

Moreover, if the suppression is both anticipatory and location-based, and not elicited by the presence of specific distractor information we would expect a P_D_ for the high-probability location even if no distractor singleton is presented there. We analyzed this component separately for when the target appeared at this location (upper right panel of Figure 2), for when the distractor singleton appeared at this location (upper middle panel of Figure 2) and for when a neutral display-element was presented at this location (no target nor distractor singleton, upper left panel of Figure 2). A two-way ANOVA with within-subjects factors *laterality* (contra-vs. ipsilateral to the high-probability location) and *type of display-element* (distractor, target, or neutral display-element) was conducted on average ERP amplitude within P_D_ time window. The results showed a main effect for laterality, *F*(1,17) = 7.96, *p* = .012, *η* _*p*_ ^2^ = .32, but not for type of display-element, *F*(2,34) = 2.15, *p* = .13, *η* _*p*_ ^2^ = .11, and no interaction was observed, *F*(2,34) = 1.93, *p* = .161, *η* _*p*_ ^2^ = .1. The activity was more positive in the contralateral side for each condition, all *t*s ≥ 2.46, all *p*s ≤ .025, indicating that a P_D_ was found regardless of which display-element was presented there. It suggests that he suppression we observed is completely spatial in origin. Note however, as can be seen in Figure 3, the mean P_D_ amplitude generated when a distractor was present at the high probability location (1.58 μV) was slightly larger than the amplitude when a neutral display-element presented at high-probability location (1.09 μV), *t* = 2.14, *p* = .047. This suggest that the presence of distracting information at this high probability location generates additional suppression above and beyond suppression that is already present when no distracting information is present. There was no reliable difference between when the target (1.43 μV) and the neutral display-element when they were presented at high-probability location, *t* = 1.2, *p* = .248.

**Figure 3.**
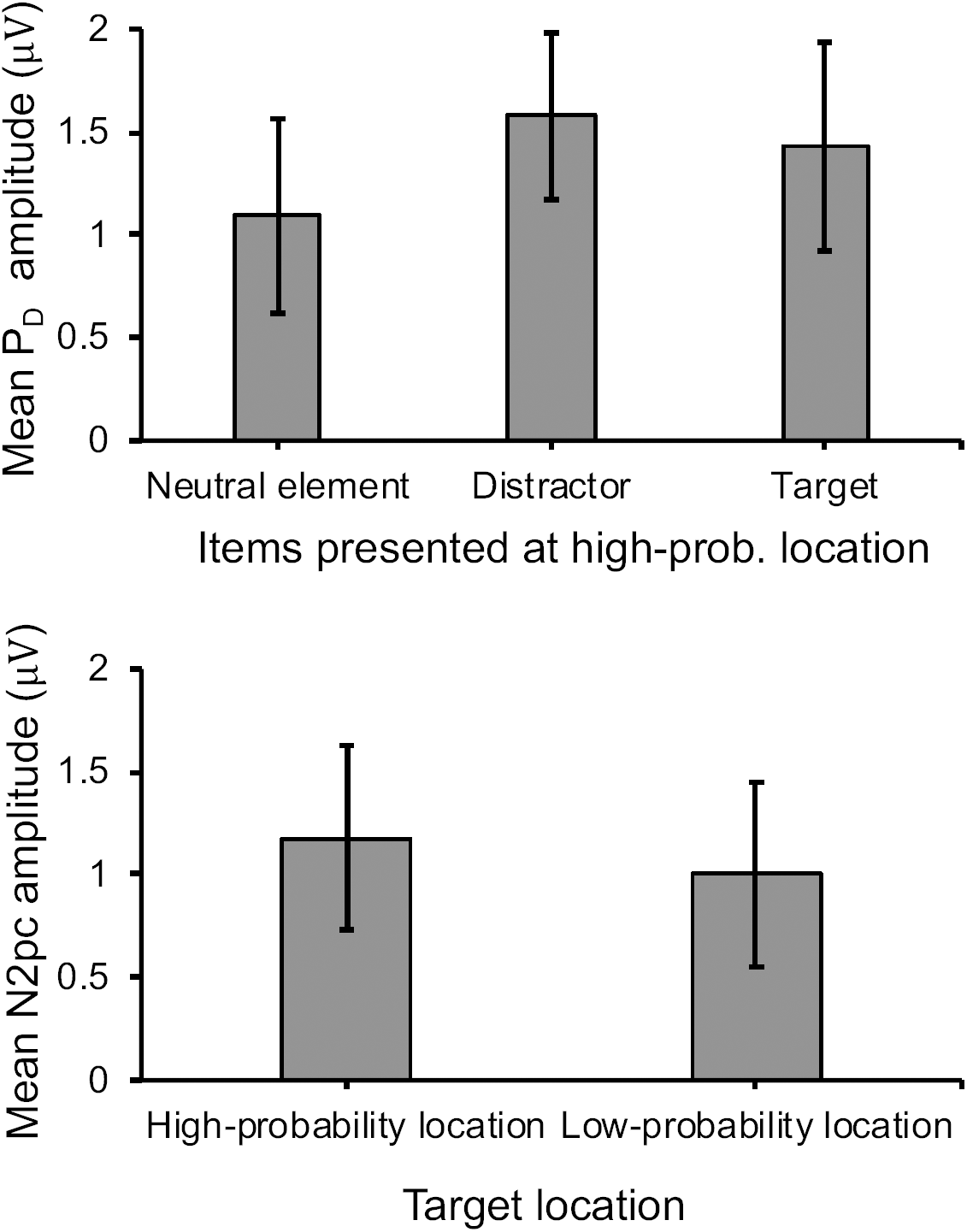
Upper panel shows mean P_D_ amplitude for different items presented at high-probability location. Lower panel shows mean N2pc amplitude for target presented at high-and low-probability locations. Error bars denote within-subjects 95% confidence intervals.

### Post-stimulus: Target-elicited N2pc

To determine whether there was attentional selection of the target and/or the distractor when presented at the low-probability location, we analyzed the N2pc towards the target or distractor singleton (lower panels of Figure 2). A two-way ANOVA with within-subjects factors *laterality* (contra vs. ipsilateral to the low-probability location) and *type of display-element* (distractor vs. target) was conducted on mean ERP amplitude within the N2pc time window (186 ms – 226 ms; see Methods for choice of time window). Only the interaction was reliable, *F*(1,17) = 7.5, *p* = .014, *η* _*p*_ ^2^ = .31. Subsequent planned comparisons tracing the source of the interaction showed a reliable N2pc only for targets, *t*(17) = 2.35, *p* = .031 (see Figure 2 lower panels).

Furthermore, we also observed what seems to be a delayed N2pc (peaked at 294 ms) when the target was presented at high-probability location, *t*(17) = 1.88, *p* = .039 (one-tailed). This amplitude of the delayed N2pc for the target at the high-probability location did not differ from the amplitude of the N2pc for the target at the low-probability location, t < 1 (see Figure 3). We used a jackknife procedure (Kiesel, Miller, Jolicœur, & Brisson, 2008) to statistically test the difference between the N2pc latencies for targets presented at the high-vs. low-probability location and obtained a reliable estimate of the numerical difference. The peak latencies between the delayed N2pc (with the average peak latency of 294 ms) for target presented at the high-probability location and the N2pc observed when the target was presented at the low-probability location (with the average peak latency of 205 ms) was significantly different, *t*(17) = 7.89, *p* < .001.

It is clear that the N2pc for targets presented at the high-probability location is weaker than in the low-probability location. If we used the same time period (± 20 ms) for calculating the N2pc as was used for targets presented at the low-probability location, then there N2pc is no longer reliable. This is probably due to the suppression (indexed by the P_D_) exerted at the high-probability location which makes the N2pc less robust. Therefore, we used the peak point to calculate the mean N2pc amplitude for targets presented at the high-probability location. When we use this method, a reliable yet less robust N2pc was observed, as we have outlined above.

### Post-stimulus: Early P_D_

Consistent with the notion of anticipatory suppression of the high-probability location, we found evidence for an early P_D_ representing an early modulation of the high-probability location in Experiment 1. A two-way ANOVA with within-subjects factors *laterality* (contra-vs. ipsilateral to the high-probability location) and *type of display-element* (distractor, target, or neutral display-element) was conducted on average ERP amplitude within the early P_D_ time window (74 ms – 114 ms; see Methods for choice of time window). The results indicated a main significant effect for laterality only, *F*(1,17) = 4.94, *p* = .04, *η* _*p*_ ^2^ = .23, and no interaction across conditions (see Figure 2), suggesting that the early P_D_ was present for the high-probability location regardless of whether a target, a distractor or a neutral display-element was presented.

### ERP results in Experiment 2 (baseline)

In the baseline experiment, the target and distractor singleton were equally likely presented at each location. Based on previous studies (Hickey et al., 2006), we expected a N2pc component for both the target and distractor singleton. A two-way ANOVA with within-subjects factors *laterality* (contra vs. ipsilateral to the singleton) and *lateralized item* (target vs. distractor) was conducted. The results showed a significant main effect for laterality, *F*(1,17) = 5.01, *p* = .039, *η* _*p*_ ^2^ = .23 (see Figure 4), but no interaction suggesting that both target and distractor singletons generated a N2pc (Hickey et al., 2006). There was indeed a significant N2pc for targets, *t*(17) = 2.17, *p* = 0.045, and a trend in the same direction for distractors, *t*(17) = 1.75, *p* = 0.099. The same time window was chosen to analyze when a neutral display-element was presented laterally (i.e., like what we did in Experiment 1). The results indicate that there was no difference between contralateral and ipsilateral sides, *t* < 1. It should be noted that unlike in Experiment 1 (which had the high-probability location), in the current baseline experiment no P_D_ was observed in any condition.

**Figure 4.**
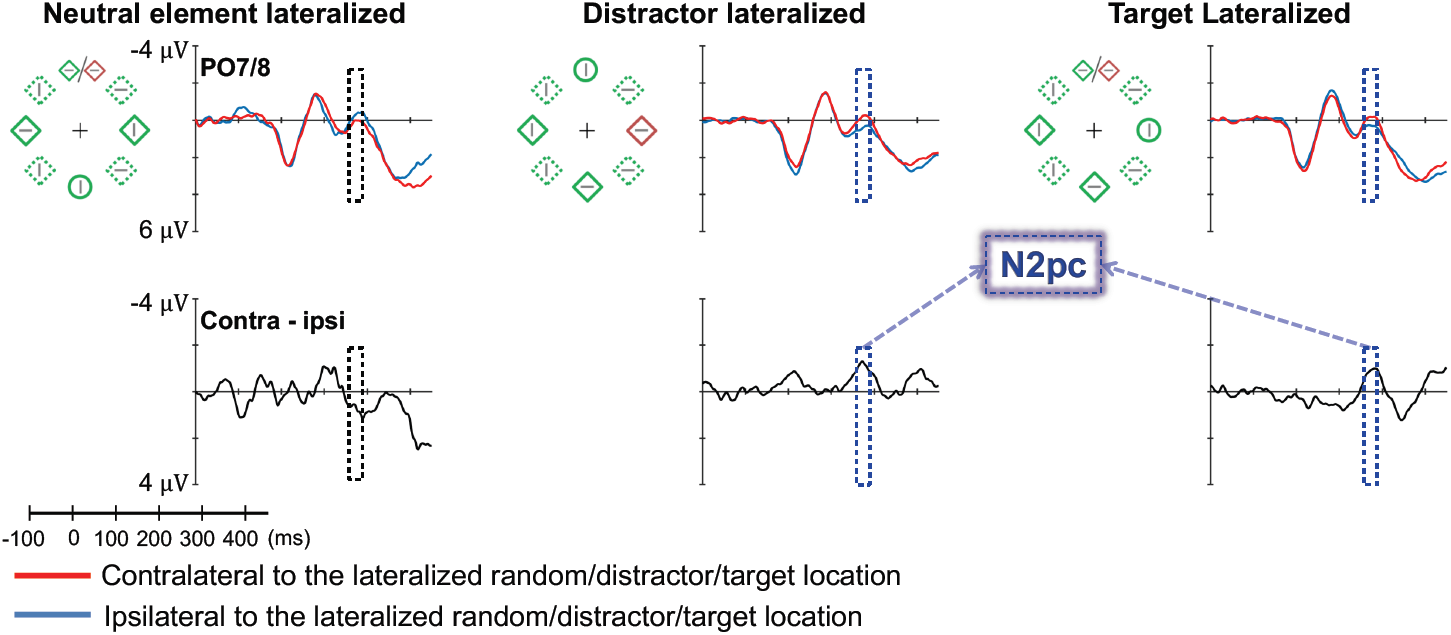
Example of the search displays in various conditions of Experiment 2. The dashed circles indicate the high-probability location. The display-elements with solid outline indicate possible target and distractor singleton locations; the display elements with dashed outline indicate locations that never contained a target or distractor singleton. Grand average ERPs recorded at PO7/PO8 and their difference waves (contra-minus ipsilateral) for different conditions were presented in different sub-panels. Blue dashed squares indicate the time window that was used to calculate mean amplitude of N2pc. All waveforms were low-pass filtered at 30 Hz (half power cutoff, 24 dB) using digital filtering.

## Discussion

Our data show that the reduction of attentional capture by a salient singleton is accompanied by pre-stimulus anticipatory suppression. Because participants have learned that the salient distractor is more likely to appear at one specific location, this location is suppressed as there was increased alpha power contralateral to this location relative to the ipsilateral location. Notably, alpha-band oscillations have been associated with neural inhibition (Jensen & Mazaheri, 2010) serving as a general gating mechanism. Previous studies have reported increased alpha power at electrodes over ipsilateral visual cortex, representing the location where participants expected distractors to appear (Kelly, Lalor, Reilly, & Foxe, 2006). This was interpreted as evidence of active attentional suppression. It has been claimed that pre-stimulus alpha operates as a gating mechanism allowing the suppression of distracting information (Moore & Armstrong, 2003).

When the salient distractor was presented at the high-probability location we found a P_D_, suggesting spatial suppression. Critically, following this P_D_ the attentional capture by salient distractor was attenuated, as no N2pc was observed towards that location. In the baseline experiment, when all locations were presented equally likely, there was no P_D_ and consequently attention was captured by the salient distractor as evidenced by an N2pc, consistent with Hickey et al (Hickey et al., 2006). Notably a P_D_ was also found when the target was presented at the high-probability location, resulting in a delayed selection (i.e., delayed N2pc) of the target consistent with the behavioral data showing delayed response time when the target is presented at that location.

Previous studies have shown that attentional suppression of distractors is typically accompanied by a P_D_ in ERP waveform (Cosman, Lowe, Woodman, & Schall, 2018; Feldmann-Wüstefeld & Vogel, 2018; Feldmann-Wüstefeld, Wykowska, & Schubö, 2013; Gaspar, Christie, Prime, Jolicœur, & McDonald, 2016; Gaspar & McDonald, 2014; Gaspelin & Luck, 2018a; Hickey et al., 2008; Sawaki & Luck, 2013). Observing a P_D_ is attributed to the processing of distracting information and is assumed to reflect active suppression of this distractor information (Burra & Kerzel, 2014; Gaspar & McDonald, 2014; Gaspelin & Luck, 2018b; Moorselaar & Slagter, 2019; Sawaki & Luck, 2013). For example, in a visual search study, Gaspelin and Luck (2018a) showed that when the salient distractor was presented, it generated a P_D_ which was correlated with behavior that showed suppression. Gaspar et al. (2016) showed that a salient distractor (e.g., a red item between green background items) generated a P_D_, while a relatively low salient distractor (e.g., an orange item between red background items) did not result in a P_D_. Overall these findings indicate that a P_D_ is only observed if a salient distractor is present, and if its salience signal needs to be actively suppressed. The findings of the current study are dramatically different in this respect. Consistent with previous finding, we do find a P_D_ when a salient distractor was present at the high-probability location (Figure 2 upper middle panel); yet, unlike previous studies we also find a P_D_ when neutral (background) elements were present (Figure 2 upper left panel); and even when the target was present at the high-probability location (Figure 2 upper right panel). In the latter case, the suppression signified by the P_D_ resulted in a delayed N2pc and a delayed behavioral response to the target.

Given the discrepancies between these findings, it is likely that the suppression that we report here is different from the suppression that is only observed when a salient distractor information is present. It is likely that this latter type of suppression is *reactive* in the sense that attention is first captured by the salient distractor (even for the briefest moment) and then is immediately suppressed (Won, Kosoyan, & Geng, 2019). Depending on the task design, it is possible that the disengagement of attention can be so fast that there are virtually very little (or no) effects to be found of the presence of the salient distractor (i.e., basically no attentional capture). It is similar to Moher and Egeth’s (2012) conception of what has been labelled as the “search and destroy” hypothesis. Moher and Egeth (2012) showed that participants that were instructed to inhibit a particular feature, could only do so after attending to the location of the to-be-ignored feature.

The type of suppression reported here can be labelled as *proactive suppression* in the sense that specific locations within the spatial priority map are suppressed (Ferrante et al., 2018; Wang & Theeuwes, 2018a, 2018b, 2018c). This suppression is spatial and feature-blind, and happens regardless of what information is present at that location (whether it is a target, a salient distractor or a neutral element). Consequently, also the neural correlates of such suppression (contralateral alpha enhancement, early P_D_, and P_D_) are observed in all these conditions. Critically, and consistent with the notion of proactive suppression, the enhanced alpha power contralateral to the high-probability location is observed prior to display onset. This indicates that the suppression is indeed proactive in the sense that the high-probability location is already suppressed before display onset. As argued, increased alpha-band oscillations are assumed to represent neural inhibition (Jensen & Mazaheri, 2010), serving as a general attentional gating mechanism.

It should be noted that P_D_ that we observed when a distractor singleton was present at the high-probability location was slightly larger than the P_D_ that we found for neutral elements presented at the high-probability location. This seems to suggest that the presence of distracting information generates additional (retroactive) suppression above and beyond the proactive suppression that was induced through statistical learning.

We also observed an early P_D_ component (80-120 ms post-stimulus) towards the high-probability location in all conditions regardless of whether a target, a salient distractor or a neutral element was presented at the high probability location. An early P_D_ (also termed positivity posterior contralateral [Ppc]; or lateralized P1) often co-occurs with the later P_D_ in displays with lateralized stimuli. Initially, Luck and colleagues (Luck, Heinze, Mangun, & Hillyard, 1990; Luck & Kappenman, 2012) found that the early P_D_ (named lateralized P1 in their study) was enhanced when the target appeared at the attended hemifield, suggesting that it may represent facilitation of early sensory processing. This is consistent with some studies suggesting that the early P_D_ is related to initial processing of stimuli or/and is assumed to represent some low level sensory imbalance between the two hemispheres. (Fortier-Gauthier, Moffat, Dell’Acqua, McDonald, & Jolicœur, 2012; Jannati, Gaspar, & McDonald, 2013),. However, in the current study, the early P_D_ was also found when non-salient neutral display elements were presented at the high-probability location, indicating that the early P_D_ does not represent early sensory processing of salience nor hemispheric imbalance, but instead represents a signature of early suppression (Weaver, van Zoest, & Hickey, 2017).

The current study shows that participants learn to suppress the location where the salient distractor is likely to appear. The pattern of results shows that relative to the high-probability location, there was pre-stimulus, contralateral alpha enhancement, an early P_D_, and the P_D._ The neural correlates were found regardless of whether a salient distractor, a salient target or a neutral element were presented at this location. This pattern of results suggests that through statistical learning the location that is likely to contain a salient distractor is proactively suppressed.

## Appendix

### Behavior results in Experiment 1

Trials on which the response times (RTs) were faster than 200 ms and slower than 2000 ms (2.17%) were excluded from analyses.

### Attentional capture effect

Mean RTs and mean error rates are presented in Figure A1A. With *distractor condition* (high-probability location, low-probability location, and no-distractor) as a factor, a repeated measures ANOVA on mean RTs showed a main effect, *F*(2, 34) = 32.09, *p* < .001, partial *η*^*2*^ = .65. Subsequent planned comparisons showed that there were significant attentional capture effects for a distractor presented on the high-probability location, *t*(17) = 4.26, *p* = .001, and the low-probability location, *t*(17) = 8.61, *p* < .001. Crucially, however, there was also a reliable difference between the high- and low-probability location, *t*(17) = 3.42, *p* = .003, suggesting that attentional capture effect was reduced for trials in which the distractor appeared at a high-probability location.

The results on error rates mimicked those for RTs. With *distractor condition* (high-probability location, low-probability location, and no-distractor) as a factor, a repeated measures ANOVA showed a main effect, *F*(2, 34) = 21.41, *p* < .001, partial *η*^*2*^ = .56. Subsequent planned comparisons showed that significant attentional capture effects were observed when a distractor singleton was presented in the high-probability location, *t*(17) = 3.82, *p* = .001, and the low-probability location, *t*(17) = 5.74, *p* < .001. Importantly, however, there was also a reliable difference between the high- and low-probability location, *t*(17) = 3.35, *p* = .004. Taken together, these results suggest that attentional capture effect was reduced for trials in which the distractor singleton appeared in a high-probability relative to the low-probability location, a straightforward replication of Wang and Theeuwes (2018a, 2018b).

### Target at the high probability distractor location

To determine whether not only attentional capture but also the efficiency of selecting the target was affected, we examined RTs in the no-distractor condition. We expected that even when the distractor was absent, observers would respond more slowly toward a target when it was presented at the high-probability distractor location relative to the low-probability location. This is exactly what we found (see the left panel of Figure A1B): Participants were slower to respond when the target was presented at the high-probability distractor location rather than at the low-probability distractor location, *t*(17) = 3.73, *p* = .002; and the response were less accuracy when the target was presented at the high-probability distractor location, *t*(17) = 2.15, *p* = .046.

**Figure A1.**
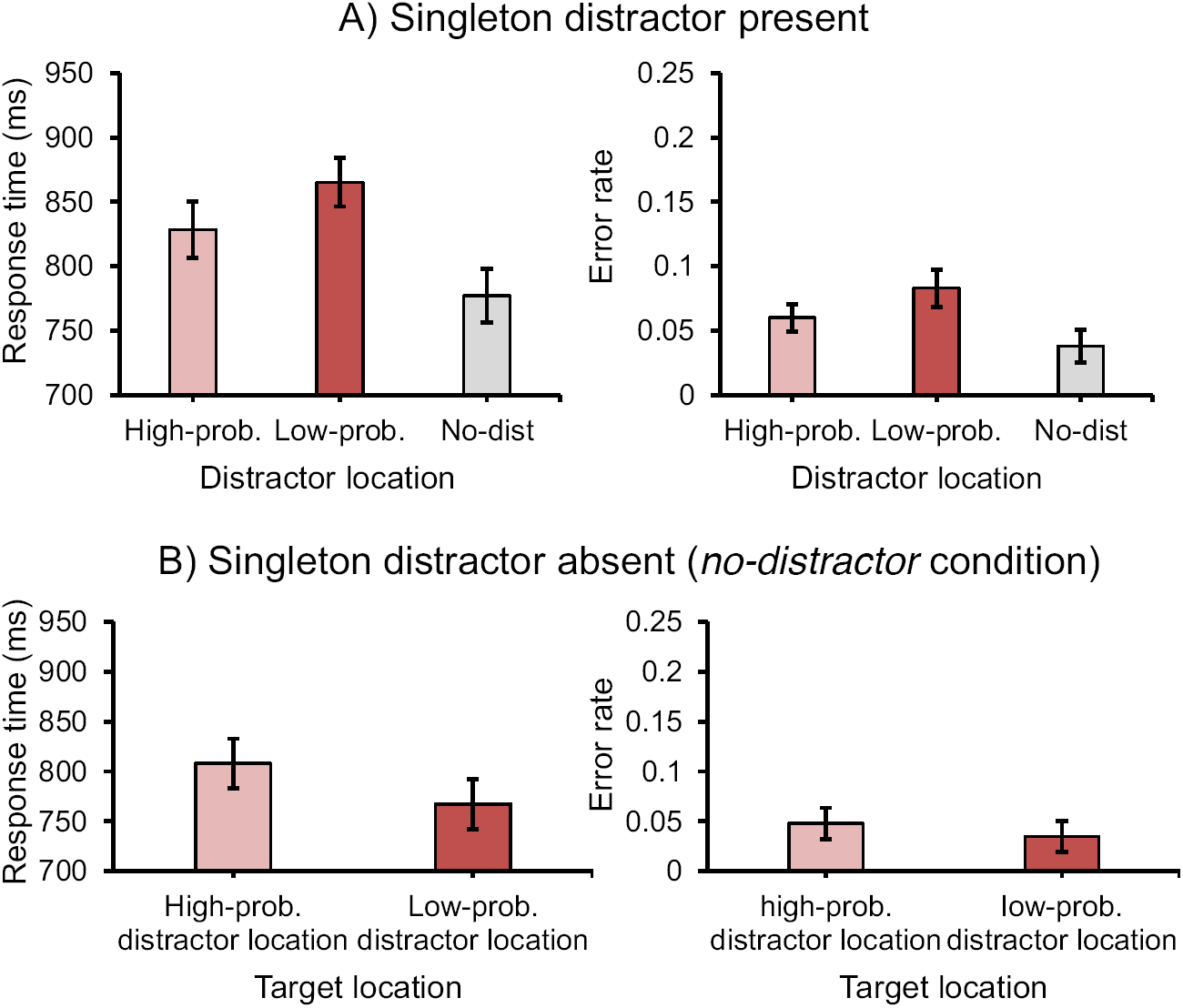
The mean response times (RTs; left panel) and the mean error rates (right panel) in singleton distractor present condition (A) and in singleton distractor absent (no-distractor) condition. Error bars denote within-subjects 95% confidence intervals.

### The spatial distribution of the suppression effect

We examined RTs and error rates of the target when the distractor singleton appeared at different locations in relation to the distance from the high-probability location. If the effect has a spatial extent, the response time (RT) and the error rate are expected to scale with distance from this area. Thus, the distractor location was divided into three distances (dist-0, dist-1, dist-2)^3^ from the high-probability location. The mean RTs and mean error rates for these conditions are presented in Figure A2A.

**Figure A2.**
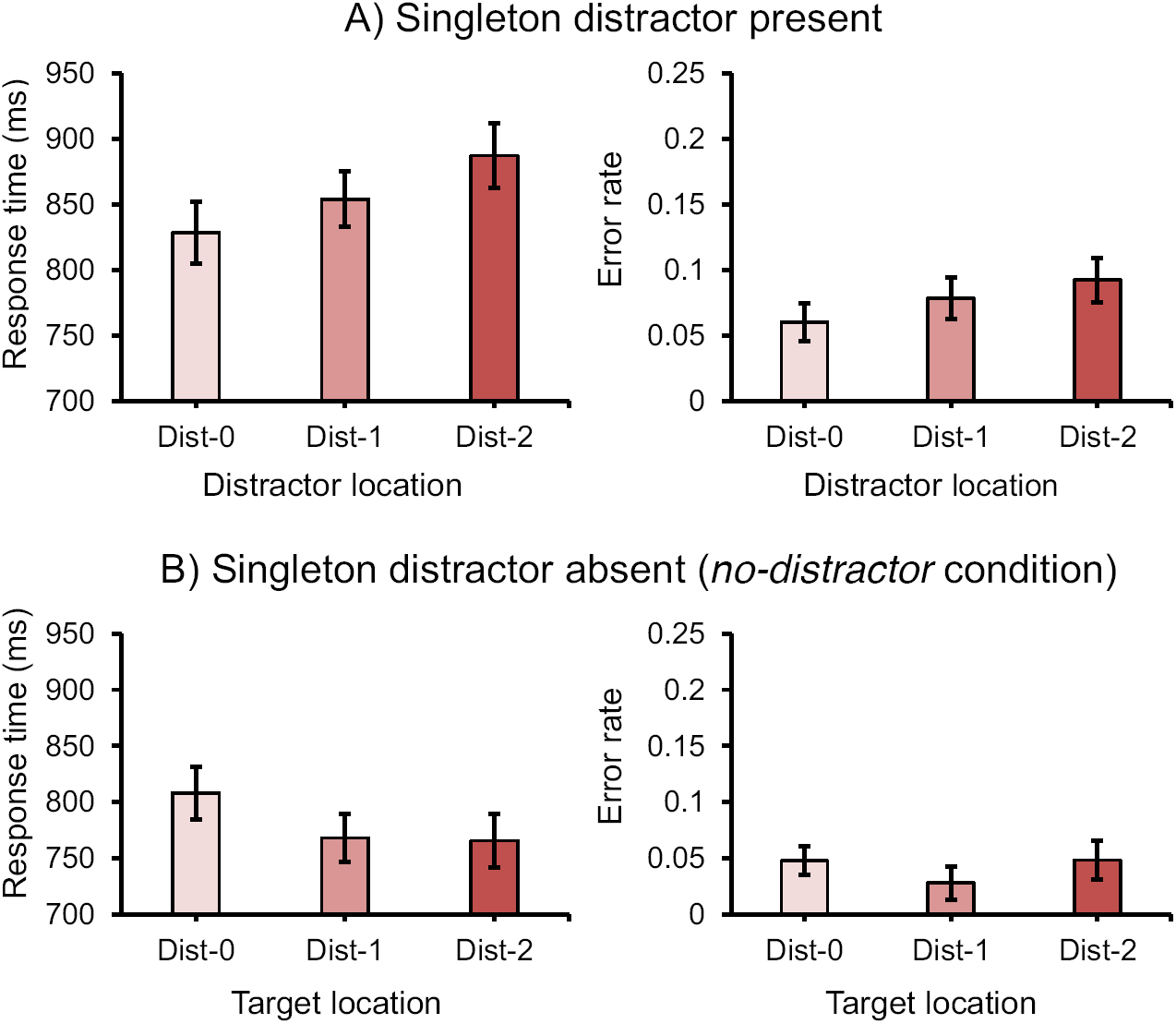
The spatial distribution of attentional capture effect by the means of response times (RTs; left panel) and error rates (right panel). Here *Dist-0* refers to the high-probability distractor location, *Dist-1* refers to the low-probability distractor location with one unit (90° polar angle) away from the high-probability distractor location, and so on. Error bars denote within-subjects 95% confidence intervals.

Repeated measures ANOVAs on mean RTs and mean error rates both showed significant main effects for *distance,F*(2, 34) = 11.57, *p* < .001, partial *η*^*2*^ = .41 and *F*(2, 34) = 7.15, *p* = .003, partial *η*^*2*^ = .3, respectively. Moreover, we fitted the data with a linear function and used their slopes to determine whether mean RTs and mean error rates changed with distance. The slopes were significantly larger than zero for mean RTs (15.4 ms per display element), *t*(17) = 4.43, *p* < .001, and mean error rates (0.8% per display element), *t*(17) = 3.73, *p* = .002, suggesting a spatial gradient (the distractor effect grew when the distractor was presented farther away from the high-probability distractor location).

In the *no-distractor* condition, there was also a significant main effect for *distance,F*(2, 34) = 7.64, *p* = .002, partial *η*^*2*^ = .31. The gradient effect was now reversed: With increased distance, the selection of the target became progressively more efficient, signified by a slope of -11.15 ms per display element that differed from zero, *t*(17) = 3.31, *p* = .004. This latter effect suggests spatial suppression that is not dependent on whether or not a distractor is present.

### Behavior results in Experiment 2

Trials on which the response times (RTs) were faster than 200 ms and slower than 2000 ms (2.24%) were excluded from analyses. Mean RTs and mean error rates are presented in Figure A3. Paired-samples t-test between different distractor conditions (*with* vs. *without distractor singleton*) showed a significant capture effect, reflected by mean RTs, *t*(17) = 10.59, *p* < .001, and mean error rates, *t*(17) = 4.16, *p* = .001.

**Figure A3.**
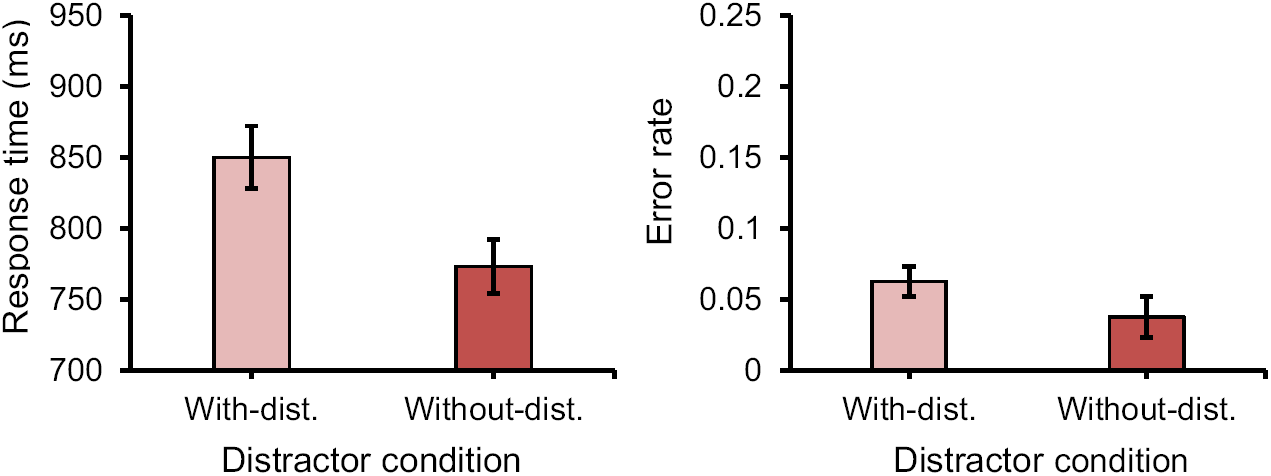
The mean response times (RTs; left panel) and the mean error rates (right panel) in different distractor condition. Error bars denote within-subjects 95% confidence intervals.

## Footnotes

1 Because in the distractor condition, the target never appears at the high probability location, it is possible that the effects reported is due to the target probability and not to the distractor probability. In a recent study we showed that the probability of the target did not matter; the effect observed was solely due to the distractor being presented more often in this one location (Failing, Wang, & Theeuwes, 2019).

2 When manually removing all the eye movement trials in Experiment 1, the results stay the same.

3 The dist-0 location refers to the high-probability distractor location; the dist-1 location means that the distractor was presented next to the high-probability distractor location (90° polar angle); the dist-2 location refers to 180° from the high-probability distractor location.

## References

Awh, E., Belopolsky, A. V., & Theeuwes, J. (2012). Top-down versus bottom-up attentional control: a failed theoretical dichotomy. Trends in Cognitive Sciences, 16(8), 437–443. doi:10.1016/j.tics.2012.06.010

Bacon, W. F., & Egeth, H. E. (1994). Overriding stimulus-driven attentional capture. Perception & Psychophysics, 55(5), 485–496. doi:10.3758/BF03205306

Burra, N., & Kerzel, D. (2014). The distractor positivity (Pd) signals lowering of attentional priority: Evidence from event-related potentials and individual differences. Psychophysiology, 51(7), 685–696. doi:10.1111/psyp.12215

Chun, M., & Jiang, Y. (1998). Contextual cueing: Implicit learning and memory of visual context guides spatial attention. Cognitive Psychology, 36(1), 28–71.

Chun, M. M., & Jiang, Y. (1999). Top-down attentional guidance based on implicit learning of visual covariation. Psychological Science, 10(4), 360–365. doi:10.1111/1467-9280.00168

Cohen, M. X. (2014). Analyzing Neural Time Series Data: Theory and Practice. MIT Press.

Cosman, J. D., Lowe, K. A., Woodman, G. F., & Schall, J. D. (2018). Prefrontal control of visual distraction. Current Biology, 28(3), 414–420.e3. doi:10.1016/j.cub.2017.12.023

Delorme, A., & Makeig, S. (2004). EEGLAB: an open source toolbox for analysis of single-trial EEG dynamics including independent component analysis. Journal of Neuroscience Methods, 134(1), 9–21. doi:10.1016/j.jneumeth.2003.10.009

van Driel, J., Gunseli, E., Meeter, M., & Olivers, C. N. L. (2017). Local and interregional alpha EEG dynamics dissociate between memory for search and memory for recognition. NeuroImage, 149, 114–128. doi:10.1016/j.neuroimage.2017.01.031

Egeth, H. E., & Yantis, S. (1997). Visual attention: Control, representation, and time course. Annual Review of Psychology, 48(1), 269–297. doi:10.1146/annurev.psych.48.1.269

Eimer, M. (1996). The N2pc component as an indicator of attentional selectivity. Electroencephalography and Clinical Neurophysiology, 99(3), 225–234. doi:10.1016/0013-4694(96)95711-9

Failing, M., & Theeuwes, J. (2017). Selection history: How reward modulates selectivity of visual attention. Psychonomic Bulletin & Review, 1–25. doi:10.3758/s13423-017-1380-y

Failing, M., Wang, B., & Theeuwes, J. (2019). Spatial suppression due to statistical regularities is driven by distractor suppression not by target activation. Attention, Perception, & Psychophysics. doi:10.3758/s13414-019-01704-9

Feldmann-Wüstefeld, T., Uengoer, M., & Schubö, A. (2015). You see what you have learned. Evidence for an interrelation of associative learning and visual selective attention. Psychophysiology, 52(11), 1483–1497. doi:10.1111/psyp.12514

Feldmann-Wüstefeld, T., & Vogel, E. K. (2018). Neural Evidence for the Contribution of Active Suppression During Working Memory Filtering. Cerebral Cortex. doi:10.1093/cercor/bhx336

Feldmann-Wüstefeld, T., Wykowska, A., & Schubö, A. (2013). Context heterogeneity has a sustained impact on attention deployment: Behavioral and electrophysiological evidence. Psychophysiology, 50(8), 722–733. doi:10.1111/psyp.12061

Ferrante, O., Patacca, A., Di Caro, V., Della Libera, C., Santandrea, E., & Chelazzi, L. (2018). Altering spatial priority maps via statistical learning of target selection and distractor filtering. Cortex, The Unconscious Guidance of Attention, 102, 67–95. doi:10.1016/j.cortex.2017.09.027

Fortier-Gauthier, U., Moffat, N., Dell’Acqua, R., McDonald, J. J., & Jolicœur, P. (2012). Contralateral cortical organisation of information in visual short-term memory: Evidence from lateralized brain activity during retrieval. Neuropsychologia, 50(8), 1748–1758. doi:10.1016/j.neuropsychologia.2012.03.032

Gaspar, J. M., Christie, G. J., Prime, D. J., Jolicœur, P., & McDonald, J. J. (2016). Inability to suppress salient distractors predicts low visual working memory capacity. Proceedings of the National Academy of Sciences, 201523471. doi:10.1073/pnas.1523471113

Gaspar, J. M., & McDonald, J. J. (2014). Suppression of salient objects prevents distraction in visual search. Journal of Neuroscience, 34(16), 5658–5666. doi:10.1523/JNEUROSCI.4161-13.2014

Gaspelin, N., & Luck, S. J. (2018a). Combined Electrophysiological and Behavioral Evidence for the Suppression of Salient Distractors. Journal of cognitive neuroscience, 30(9), 1265–1280. doi:10.1162/jocn_a_01279

Gaspelin, N., & Luck, S. J. (2018b). The role of inhibition in avoiding distraction by salient stimuli. Trends in Cognitive Sciences, 22(1), 79–92. doi:10.1016/j.tics.2017.11.001

Groppe, D. M., Makeig, S., & Kutas, M. (2009). Identifying reliable independent components via split-half comparisons. NeuroImage, 45(4), 1199–1211. doi:10.1016/j.neuroimage.2008.12.038

Hickey, C., Di Lollo, V., & McDonald, J. J. (2008). Electrophysiological indices of target and distractor processing in visual search. Journal of Cognitive Neuroscience, 21(4), 760–775. doi:10.1162/jocn.2009.21039

Hickey, C., McDonald, J. J., & Theeuwes, J. (2006). Electrophysiological Evidence of the Capture of Visual Attention. Journal of Cognitive Neuroscience, 18(4), 604–613. doi:10.1162/jocn.2006.18.4.604

Itti, L., & Koch, C. (2001). Computational modelling of visual attention. Nature Reviews Neuroscience, 2(3), 194–203. doi:10.1038/35058500

Jannati, A., Gaspar, J. M., & McDonald, J. J. (2013). Tracking target and distractor processing in fixed-feature visual search: Evidence from human electrophysiology. Journal of Experimental Psychology: Human Perception and Performance, 39(6), 1713.

Jensen, O., & Mazaheri, A. (2010). Shaping functional architecture by oscillatory alpha activity: gating by inhibition. Frontiers in Human Neuroscience, 4. doi:10.3389/fnhum.2010.00186

Jiang, Y., & Chun, M. M. (2001). Selective attention modulates implicit learning. The Quarterly Journal of Experimental Psychology Section A, 54(4), 1105–1124. doi:10.1080/713756001

Kayser, J., & Tenke, C. E. (2015). Issues and considerations for using the scalp surface Laplacian in EEG/ERP research: A tutorial review. International Journal of Psychophysiology, On the benefits of using surface Laplacian (current source density) methodology in electrophysiology, 97(3), 189–209. doi:10.1016/j.ijpsycho.2015.04.012

Kelly, S. P., Lalor, E. C., Reilly, R. B., & Foxe, J. J. (2006). Increases in alpha oscillatory power reflect an active retinotopic mechanism for distracter suppression during sustained visuospatial attention. Journal of Neurophysiology, 95(6), 3844–3851. doi:10.1152/jn.01234.2005

Kiesel, A., Miller, J., Jolicœur, P., & Brisson, B. (2008). Measurement of ERP latency differences: A comparison of single-participant and jackknife-based scoring methods. Psychophysiology, 45(2), 250–274. doi:10.1111/j.1469-8986.2007.00618.x

Klimesch, W. (1996). Memory processes, brain oscillations and EEG synchronization. International Journal of Psychophysiology, New Advances in EEG and cognition, 24(1), 61–100. doi:10.1016/S0167-8760(96)00057-8

Klimesch, W. (1999). EEG alpha and theta oscillations reflect cognitive and memory performance: a review and analysis. Brain Research Reviews, 29(2), 169–195. doi:10.1016/S0165-0173(98)00056-3

Klimesch, W. (2012). Alpha-band oscillations, attention, and controlled access to stored information. Trends in Cognitive Sciences, 16(12), 606–617. doi:10.1016/j.tics.2012.10.007

Lopez-Calderon, J., & Luck, S. J. (2014). ERPLAB: an open-source toolbox for the analysis of event-related potentials. Frontiers in Human Neuroscience, 8. doi:10.3389/fnhum.2014.00213

Luck, S. J., Heinze, H. J., Mangun, G. R., & Hillyard, S. A. (1990). Visual event-related potentials index focused attention within bilateral stimulus arrays. II. Functional dissociation of P1 and N1 components. Electroencephalography and Clinical Neurophysiology, 75(6), 528–542. doi:10.1016/0013-4694(90)90139-B

Luck, S. J., & Kappenman, E. S. (2012). ERP components and selective attention. The Oxford handbook of event-related potential components, 295–327.

Maris, E., & Oostenveld, R. (2007). Nonparametric statistical testing of EEG- and MEG-data. Journal of Neuroscience Methods, 164(1), 177–190. doi:10.1016/j.jneumeth.2007.03.024

Moore, T., & Armstrong, K. M. (2003). Selective gating of visual signals by microstimulation of frontal cortex. Nature, 421(6921), 370–373. doi:10.1038/nature01341

Moorselaar, D. van, & Slagter, H. A. (2019). Learning what is irrelevant or relevant: Expectations facilitate distractor inhibition and target facilitation through distinct neural mechanisms. bioRxiv, 565069. doi:10.1101/565069

Oostenveld, R., Fries, P., Maris, E., & Schoffelen, J.-M. (2011). FieldTrip: Open Source Software for Advanced Analysis of MEG, EEG, and Invasive Electrophysiological Data. Intell. Neuroscience, 2011, 1:1–1:9. doi:10.1155/2011/156869

Perrin, F., Pernier, J., Bertrand, O., & Echallier, J. F. (1989). Spherical splines for scalp potential and current density mapping. Electroencephalography and Clinical Neurophysiology, 72(2), 184–187. doi:10.1016/0013-4694(89)90180-6

Sawaki, R., & Luck, S. J. (2010). Capture versus suppression of attention by salient singletons: Electrophysiological evidence for an automatic attend-to-me signal. Attention, Perception, & Psychophysics, 72(6), 1455–1470. doi:10.3758/APP.72.6.1455

Sawaki, R., & Luck, S. J. (2013). Active suppression after involuntary capture of attention. Psychonomic bulletin & review, 20(2), 296–301. doi:10.3758/s13423-012-0353-4

Theeuwes, J. (1991). Cross-dimensional perceptual selectivity. Perception & Psychophysics, 50(2), 184–193. doi:10.3758/BF03212219

Theeuwes, J. (1992). Perceptual selectivity for color and form. Perception & Psychophysics, 51(6), 599–606. doi:10.3758/BF03211656

Theeuwes, J. (2010). Top–down and bottom–up control of visual selection. Acta Psychologica, 135(2), 77–99. doi:10.1016/j.actpsy.2010.02.006

Theeuwes, J. (2018). Visual selection: usually fast and automatic; seldom slow and volitional. Journal of Cognition, 1(1). doi:10.5334/joc.13

Theeuwes, J. (2019). Goal-driven, stimulus-driven, and history-driven selection. Current Opinion in Psychology, Attention & Perception, 29, 97–101. doi:10.1016/j.copsyc.2018.12.024

Vries, I. E. J. de, van Driel, J., & Olivers, C. N. L. (2017). Posterior α EEG Dynamics Dissociate Current from Future Goals in Working Memory-Guided Visual Search. Journal of Neuroscience, 37(6), 1591–1603. doi:10.1523/JNEUROSCI.2945-16.2016

Wang, B., & Theeuwes, J. (2018a). Statistical regularities modulate attentional capture. Journal of Experimental Psychology: Human Perception and Performance, 44(1), 13–17. doi:10.1037/xhp0000472

Wang, B., & Theeuwes, J. (2018b). How to inhibit a distractor location? Statistical learning versus active, top-down suppression. Attention, Perception, & Psychophysics, 1–11. doi:10.3758/s13414-018-1493-z

Wang, B., & Theeuwes, J. (2018c). Statistical regularities modulate attentional capture independent of search strategy. Attention, Perception, & Psychophysics, 1–12. doi:10.3758/s13414-018-1562-3

Weaver, M. D., van Zoest, W., & Hickey, C. (2017). A temporal dependency account of attentional inhibition in oculomotor control. NeuroImage, 147, 880–894. doi:10.1016/j.neuroimage.2016.11.004

Won, B.-Y., Kosoyan, M., & Geng, J. J. (2019). Evidence for second-order singleton suppression based on probabilistic expectations. Journal of Experimental Psychology: Human Perception and Performance, 45(1), 125.

Worden, M. S., Foxe, J. J., Wang, N., & Simpson, G. V. (2000). Anticipatory biasing of visuospatial attention indexed by retinotopically specific alpha-band electroencephalography increases over occipital cortex. Journal of Neuroscience, 63.

